# Differential neuronal vulnerability to C9orf72 repeat expansion driven by Xbp1 transcription signature

**DOI:** 10.1101/2023.11.20.567861

**Authors:** Dunxin Shen, Alec Vincent, Evan Udine, Yazead Buhidma, Sharifah Anoar, Elli Tsintzas, Marie Maeland, Rosa Rademakers, Adrian M Isaacs, Carlo Frigerio, Marka van Blitterswijk, Tammaryn Lashley, Teresa Niccoli

## Abstract

A G4C2 repeat expansion in the gene *C9orf72* (C9) is the most common genetic cause of sporadic and familial frontotemporal dementia (FTD) and amyotrophic lateral sclerosis (ALS). What determines why cell death is triggered only in specific neuronal populations, while others remain ‘protected’ or are less susceptible to disease is still an open question. In particular, whether it is the transcriptional response to the accumulation of toxic insults or the initial cellular state that determines their vulnerability is still unknown.

We have carried out a large-scale profiling of single cell transcriptional signatures throughout disease development in a *Drosophila* model of C9 repeat toxicity. This enabled us to monitor transcriptional shifts and track changes in cell populations during disease progression. We have identified neuronal populations which are depleted in response to C9 repeat expression, and therefore vulnerable to toxicity. On the other hand, other neuron types are resistant to toxicity, and maintain their cell number during disease progression. Our findings suggest that a major determinant of vulnerability is the transcriptional state of the cell before it is exposed to C9 repeat expression. We have identified a conserved transcriptional profile that is associated with resistance to C9 repeat toxicity. Neurons resistant to disease display a higher expression of genes involved in protein homeostasis, with Xbp1 identified as a crucial transcription factor determining neuronal vulnerability.

## Introduction

Dementia, the most common neurodegenerative disease, is currently one of the leading causes of death in industrialized countries, affecting nearly 50 million people worldwide [1]. Concerted research efforts have successfully identified many of the causative agents of dementia: mis-processed Aß peptides triggers Alzheimer’s disease [2], and alpha-synuclein accumulation leads to Parkinson’s disease. A hexanucleotide repeat expansion in the *C9orf72* gene (C9) has been identified as the most common genetic cause of FTD/ALS (frontotemporal dementia/amyotrophic lateral sclerosis), two diseases with distinct clinical presentations and where distinct neuronal populations are affected, but which share pathological hallmarks and genetic causes, and are considered two extremes of the same disease spectrum [3, 4].

The molecular cascades leading to neuronal cell death remain unknown and a major, outstanding question is why cell death is triggered only in specific neuronal populations, whereas others remain ‘protected’ [5]. This neuronal type-specific vulnerability to cell death is linked to the toxic element. Aß accumulation in Alzheimer’s disease causes hippocampal neuron death, whereas *PRKN* mutations in Parkinson’s disease lead to the degeneration of nigral neurons [5], yet the genes causal for these diseases are expressed throughout the brain. The intrinsic biological properties of neurons have been proposed as a reason for selective vulnerability of neuronal populations to different toxic insults [6]. It has been suggested that different types of neurons exhibit varying susceptibility to the disruption of specific cellular protein homeostasis networks, which may contribute to their vulnerability to specific insults which lead to neurodegenerative diseases. This susceptibility is disease-specific, as indicated by genome-wide association studies (GWAS) linking disease-associated genes to distinct proteostasis pathways: the ubiquitin proteasome pathways in FTD/ALS as opposed to the lysosome in pure FTD and Dementia with Lewy bodies [7].

To understand how accumulation of toxic insults can lead to neurodegeneration we focused on C9, which causes death of cortical neurons in FTD and motor neurons in ALS. C9 is a dominant, highly penetrant, genetic mutation that causes age-dependant neurodegeneration [8]. The C9 expansion is composed by the repetition of a G4C2 repeat, which is generally found in less than 30 copies in the general population and can expand to thousands of copies in disease. These repeats are transcribed both in the sense and antisense direction to generate highly stable RNA species which can be translated in a non-canonical form of translation, called repeat associated non-ATG mediated translation (RAN) [9], to generate 5 different Dipeptide Repeat Proteins (DPRs), all of which are found in patients’ brains [10]. We previously developed *Drosophila* models of C9, which demonstrated for the first time that C9 repeats are highly toxic to adult neurons and that the toxicity is mostly associated to the increase in DPR proteins [11].

We used this model to identify which neuronal populations are vulnerable and resistant to disease and to uncover transcriptional signatures associated with toxicity. We have carried out single-cell RNA sequencing of *Drosophila* brains across a time-course of induction of the expression of C9 repeat expansion. We have identified neuronal populations which are resistant and vulnerable to disease, and found pathways associated with both. In particular we find a distinct baseline signature, associated with protein homeostasis, which modulates the vulnerability of neurons to C9 toxicity, suggesting it is not how a neuron responds but how it its initial cellular environment which determines its vulnerability.

## Results

### Single-cell RNA sequencing of the C9 repeat expressing fly brains identifies over 200 cell types

In order to understand how different cellular populations respond to toxic insults associated with dementia, we monitored, at single-cell resolution, the transcriptional response to C9 repeat expression. In our model, 36 G4C2 repeats are expressed via the neuronal driver elavGS. The activation of this driver is triggered by the drug RU486 (RU), given to the flies after they eclose. This ensures that the repeats are expressed exclusively in adult neurons. C9 expressing flies display a very rapid onset phenotype and start dying within 10 days of induction (Figure S1A)[11]. We decided to analyse the transcriptional response throughout disease development, until the population started declining. We placed flies onto RU containing food every 2 days to induce C9 repeat expression, to generate 5 groups of flies which were exposed to C9 repeat expression for 0, 2, 4, 6 and 8 days separately, all flies were the same age (Figure S1B), thus eliminating any confounding ageing component from the analysis. Brains from all groups were dissected on the same day and the single-cell suspensions were prepared by digestion with dispase and collagenase, following by passage through a 10 μm cell strainer. We ensured cell viability was over 90% and then sequenced single-cell cDNA libraries from each sample (Figure S1C). The obtained reads were aligned with the fly genome, and we filtered out cells with less than 200 genes and genes detected less than 3 cells. We also filtered out cells with more than 3000 genes and cells with mitochondrial DNA higher than 5%. The cells retained were taken for further analysis [12].

Following quality control filtering, we retained 101,806 high-quality cells which are evenly distributed over 3 experimental batches when projected onto uniform manifold approximation and projection (UMAP, Figure 1A). They were, however, unevenly distributed over the 5 time points in a few areas when projected onto UMAP (Figure 1B), indicating C9 repeats having an impact on the fly brain transcriptome. We checked the expression of *nSyb* and *repo*, which showed our sampled brain cells consisted of 90.7% neuronal cells, 2.5% glial cells, and 6.8% other types of cells (Figure 2C, 2D, S2B-D), in keeping with other single cell analyses of fly brains [13, 14]. Based on their transcriptomic signature, cells were assigned to 214 clusters, using Leiden in Scanpy at a resolution of 10.0 (Figure 2E) [15, 16], this is consistent with previous clustering analysis of fly brain single cell sequencing data [17]. The neuronal clusters were further identified, based on the expression of key marker genes, such as cholinergic (*VAChT* or *ChAT*), glutamatergic (*VGlut)*, GABAergic (*Gad1*), serotoninergic (*SerT*), dopaminergic (*ple*), and octopaminergic/tyraminergic (*Tdc2*) neurons based on their expression of marker genes (Figure 1F). The cell numbers in each neurotransmitter category do not change across the time points, indicating there is no selective depletion of a population associated with a particular neurotransmitter (Figure S2E-F).

**Figure 1.**
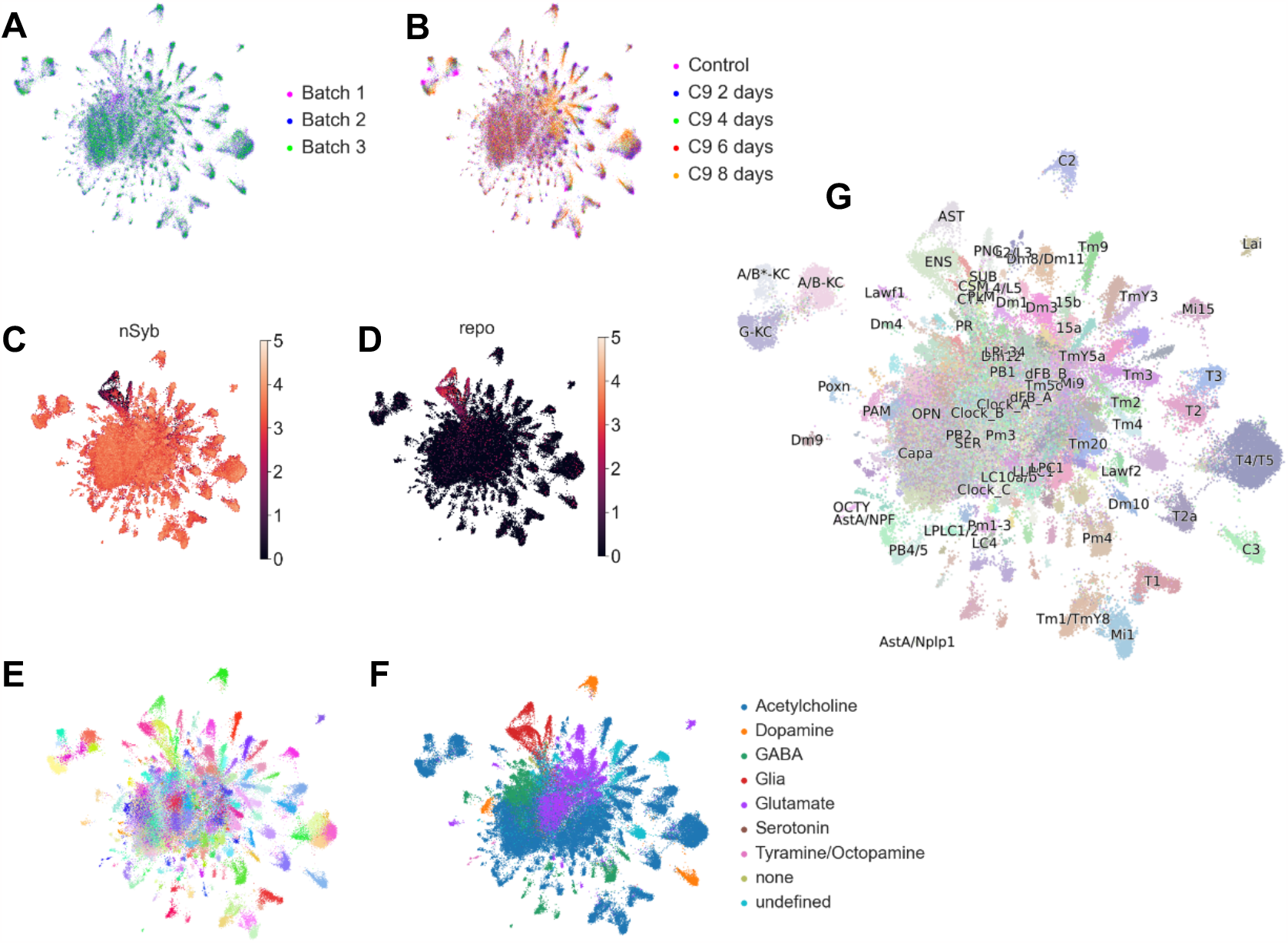
C9 Fly Brain Cells of 3 Experimental Batches across 5 Time Points Projected onto UMAP with Different Types of Labels. (A) Cells are evenly distributed over 3 experimental batches on the UMAP. (B) Cells are unevenly distributed across 5 times points in some areas on the UMAP. (C) Normalised and log-transformed *nSyb* expression levels indicating neuronal cells on the UMAP. (D) Normalised and log-transformed *repo* expression levels indicating glial cells on the UMAP. (E) Cells divided into 214 clusters using Leiden with resolution at 10.0 on the UMAP. (F) Neuronal cells divided into categories of different neurotransmitter; “none” refers to cells containing no marker genes of neurotransmitter; “undefined” refers to cells containing more than one marker genes of neurotransmitter. (G) Cells with transferred labels using 70% threshold from the published fly brain single-cell atlas; AST, astrocyte-like glia; C, centrifugal neurons; CSM, chiasm glia; CTX, cortex glia; ENG, ensheathing glia; GMC, ganglion mother cells; L, lamina monopolar neurons; LAM, lamina neurons; LPC, lamina precursor cells; NB, neuroblasts; ONE, OL neuroepithelium; PB, protocerebral bridge neurons; PLM, plasmatocytes; PNG, perineurial glia; PR, photoreceptors; PXN, poxn-neurons of ellipsoid body; SUB, sub-perineurial glia; SUR, surface glia; Tm, transmedullary neurons.

**Figure 2.**
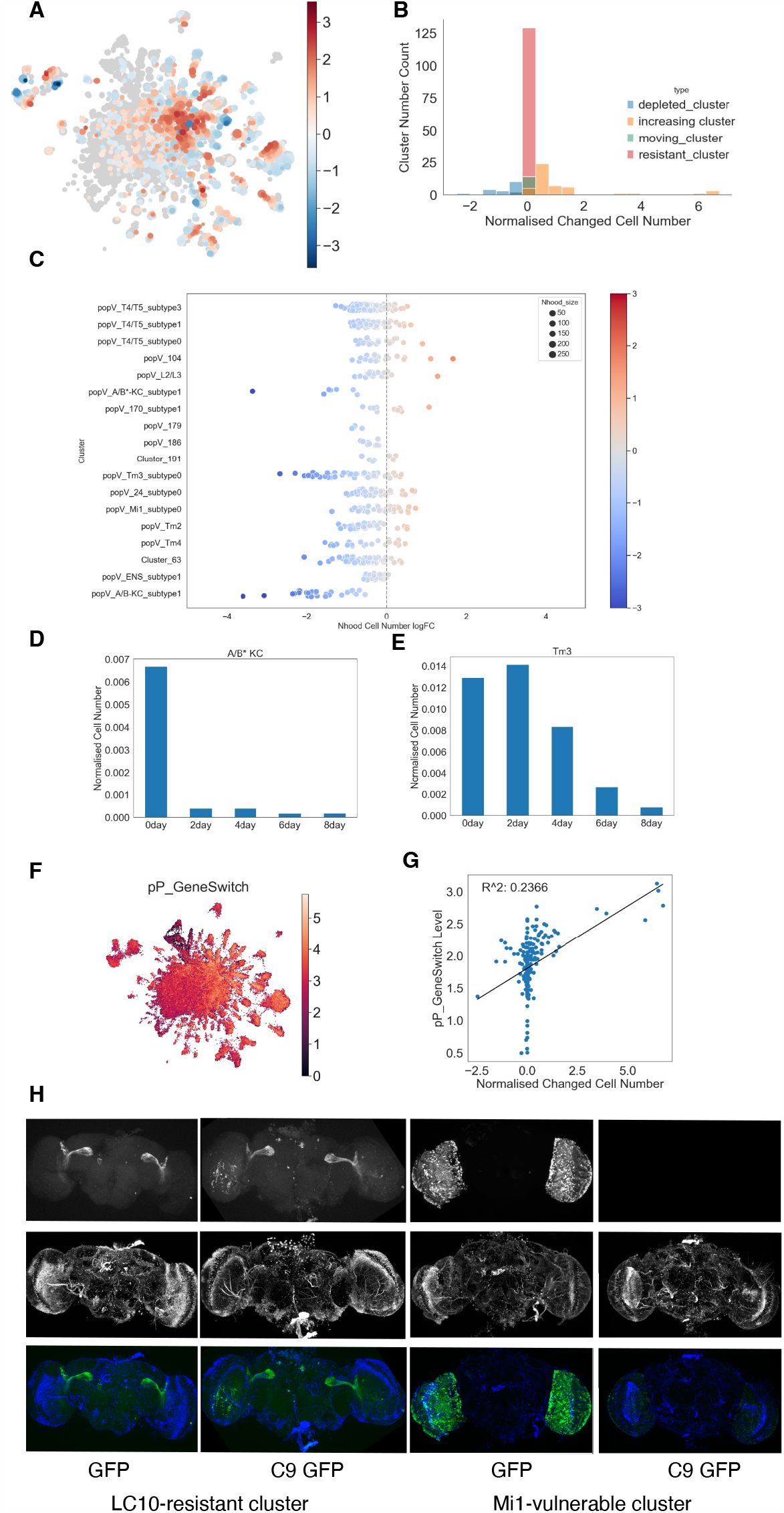
Differential Abundance Analysis with Milo Revealing Vulnerable and Resistant Cells. (A) Milo calculated neighbourhoods on the UMAP. Blue dots refer to neighbourhoods with decreased cell number; Red dots refer to neighbourhoods with increased cell number; grey dots refer to neighbourhoods without significant cell number change. (B) Bin diagram showing the number of clusters displaying a given normalised cell number change. Resistant clusters (showing 0 cell number change) were the majority. (C) Swarm plot of all Milo calculated neighbourhoods in each depleted neuronal cluster. (D) Bar plot of normalised cell numbers in A/B* KC cells across 5 time points. (E) Bar plot of normalised cell numbers in Tm3 cells across 5 time points. (F) Normalised and log-transformed expression levels of the transgene GeneSwitch. (G) Dot plot and linear regression of every cluster’s normalised cell number change and average expression level of GeneSwitch. (H) Fluorescent imaging of fly brains with membrane conjugated GFP expressed in resistant LC10 neurons and vulnerable Mi1 neurons with and without to C9 expression (Green: GFP; Blue: DAPI)

To allocate neuronal identities to our clusters, we transferred the labels of cell types from a published fly brain single-cell atlas to our dataset using a label-transferring algorithm called PopularVote (popV, Figure 1G) [18]. There was good agreement between the label-transferred cell type and our Leiden clustering. 127 out of 214 clusters were assigned a transferred label when setting the threshold of cells within a cluster sharing the same label to 70% (Table S1).

### C9 expression leads to selective neuronal depletion

To investigate whether any populations were selectively depleted in response to C9 expression we used the Milo tool, which re-assigns all cells to smaller, highly related clusters, called neighbourhoods, and then calculates changes in cell number in these neighbourhoods across the timepoints (Figure 2A) [19]. We then re-allocated each neighbourhood back to its corresponding cluster and calculated a total normalised cell number change across the timepoints to determine if one cluster is depleted or stable. (See Methods for details). Depending on the cell number changes, all the clusters were divided into four categories – clusters with overall cell number decreasing more than 20% as depleted clusters, clusters with normalised cell number increasing more than 20% as increasing clusters, the rest of clusters as resistant clusters. Within some resistant clusters, we noticed some neighbourhoods increasing and others decreasing, if more than 20% of neighbourhoods within a cluster were found to both increase and 20% decrease, we classified this as a “moving” cluster. We presume the transcription profiles in these particular clusters changed substantially due to the C9 toxicity but that this did not lead to overt cellular depletion (Figure 2B). With this analysis we identified 18 clusters that were selectively depleted across the timepoints, and were therefore vulnerable to C9 toxicity, and 131 which maintained their cell numbers across the timepoints and were therefore resistant to toxicity. The vulnerable clusters included Kanyon Cell (KC) neurons, which are part of the mushroom body and are involved in olfactory learning and memory, and a number of clusters (Tm2, Tm3, Tm4, Mi1, T4/T5, L2/L3) located in the medulla, a part of the visual processing centre of the fly brain.

This analysis indicates that, within a *Drosophila* brain, different cell populations are displaying differential vulnerability to C9 expression. Most of vulnerable clusters are neuronal except one which corresponds to ensheathing glia. There are 5 glial clusters and one photoreceptor cluster in the resistant clusters, with all the others being neuronal. Interestingly, depleted clusters showed different patterns of depletion, for example, A/B* KC cluster presents a fast depletion pattern with cells decreasing dramatically from 2 days after C9 repeats exposure (Figure 2D), while Tm3 cells gradually decreased from 4 days after C9 repeats exposure (Figure 2E), suggesting that the dynamics of depletion are also cell-type specific.

It is possible that the reason cells are being depleted is simply because of higher expression of the C9 repeats in those cells. We could not, for technical reasons, identify C9 repeats in our dataset, we therefore checked the expression of the driver, Gal4-Geneswitch, in our dataset (Figure 2F). We found no positive correlation between each cluster’s cell number change and the expression level of the Gal4-GS (Figure 2G, R^2^ > 0.05), indicating that the cells being depleted do not express higher levels of the driver, and by extension, the repeats.

To validate the depleted and resistant clusters identified in our single-cell analysis, we used driver lines expressed exclusively in resistant cluster LC10 and vulnerable clusters Tm2, Tm4, T4/T5 and Mi1 [20-22]. We expressed the C9 repeats, specifically in these cell types together with mCD8-GFP and compared the GFP signals to controls just expressing GFP via fluorescent imaging of fixed brains. The GFP signals were reduced in the fly brains where C9 repeats were expressed in the depleted clusters compared to non-C9 control, while the GFP signals maintained at the same level when C9 repeats were expressed in LC10 neurons compared to non-C9 controls (Figure 2H and S3A-C).This confirms that the clusters identified by bioinformatics analysis as vulnerable are depleted in the presence of C9 repeats whereas resistant clusters are not.

### Vulnerable and resistant neurons display common and distinct signatures in response to C9 repeat expression

As we are particularly interested in neuronal selective vulnerability, we concentrated our analysis going forward on the 17 depleted neuronal clusters (Figure 2C and Table S2) and the 126 resistant neuronal clusters (Figure S3D and Table S3).

To determine whether differential vulnerability is determined by how different neuronal populations respond to the expression of the repeats, we analysed the transcriptional response of clusters to the expression of C9 repeats. We performed differential gene expression (DE) analysis using a Wilcoxon rank test on each cluster in turn across all timepoints, comparing time point 0 to all other timepoints in turn (0 and 2, 0 and 4, 0 and 6, and 0 and 8) and combined any significantly differentially expressed genes (adjusted p-value < 0.05) to generate a comprehensive list of DE genes for each cluster. We then ran GO analysis separately on downregulated genes and upregulated genes (we termed this C9 progression response) [23, 24]. We identified 4 categories for C9 progression: pathways upregulated in depleted clusters, pathways downregulated in depleted clusters, pathways upregulated in resistant clusters, and pathways downregulated in resistant clusters (Table S4). By comparing the pathways in the 4 categories in C9 progression, we found that pathways associated with ribosomes and translation, which have been extensively associated with C9 toxicity [25-28] are commonly downregulated in both vulnerable and resistant neurons (Figure 3A, circled in magenta, Figure 3B-C). The strongest unique signatures were pathways downregulated either specifically in resistant clusters (Figure S4) and pathways which are uniquely downregulated in depleted clusters (Fig 3A, circled in blue, Figure 3D). Strikingly, pathways downregulated in depleted clusters were related to ribosomal assembly and, mostly, to mitochondria’s electron transport chain (Figure 3D, highlighted in red), suggesting there is a breakdown in mitochondrial respiration in vulnerable neurons.

**Figure 3.**
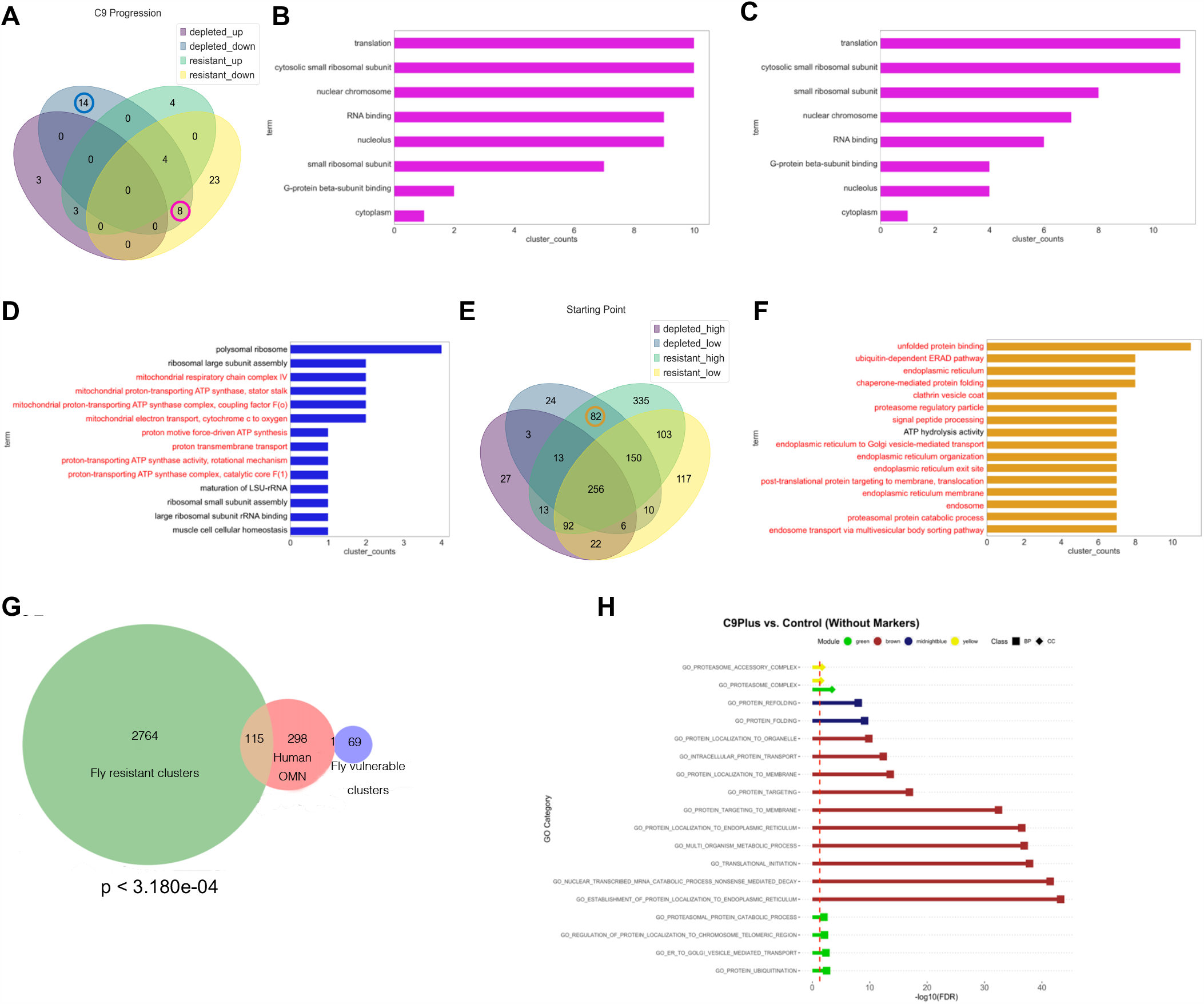
Differential Gene Expression Analysis and Gene Ontology Analysis. (A) Venn diagram of number of pathways associated with depleted and resistant neurons in C9 Progression. (B) Bar plot of top pathways downregulated in both depleted and resistant neuronal clusters, showing in how many depleted clusters these were found. (C) Bar plot of top pathways downregulated in both depleted and resistant neuronal clusters, showing in how many resistant clusters these were found. (D) Bar plot of top pathways uniquely downregulated in depleted neuronal clusters. (E) Venn diagram of number of pathways associated with depleted and resistant neurons in Starting Point analysis. (F) Bar plot of high-frequent pathways highly expressed in resistant clusters, and lowly expressed in depleted clusters. (G) Venn diagram of genes highly expressed in at least 5 resistant (green) and 5 vulnerable (blue) clusters in *Drosophila* and genes upregulated in human oculomotor neurons (OMN) . The overlap between genes highly expressed in resistant neurons in *Drosophila* and genes upregulated in human oculomotor neurons is significant (Hypergeometric test, p < p < 3.180e-04). (H) WGCNA Pathway Enrichment results for disease associated modules in the C9Plus vs. Control comparison. The dotted red line represents and FDR of 0.05. Class BP refers to GO:Biological Process and CC refers to GO:Cellular Component.

### Protein homeostasis networks modulate C9 selective vulnerability

The analysis of neuronal response to C9 repeat expression highlighted a few pathways, but did not provide a prominent differential signature related to neuronal vulnerability, we wondered whether, instead of how neurons respond to toxic insults, neuronal vulnerability could be mostly determined by the initial cellular environment of the neuron. To test this, we analysed the basal transcriptional state of neuronal populations (starting point). We performed DE analysis using a Wilcoxon rank test on the time 0 transcriptome of each cluster against all others and combined all genes DE from vulnerable, and resistant clusters. We then ran GO analysis using GOATOOLS for upregulated and downregulated genes separately (Table S5), identifying 4 categories of GO pathways: pathways upregulated in depleted clusters, pathways downregulated in depleted clusters, pathways upregulated in resistant clusters, and pathways downregulated in resistant clusters. We were interested in identifying GO pathways that were highly expressed in resistant clusters, and downregulated in depleted clusters, as these could potentially be modulators of vulnerability. Overlapping the pathways in the 4 starting point categories, found 82 such GO pathways (Figure 3E, circled in orange, Figure 3F). Interestingly, most of them are highly related to cellular protein homeostasis (Figure 3F, highlighted in red), indicating that neuronal populations possessing higher capability of proteostasis might have higher resistance against C9 repeat toxicity.

To check whether the starting point transcriptional signature of resistant neurons is conserved we looked for commonalities between our signature and that of ocular motor neurons, which are known to be resistant to ALS [29]. Allodi, et al. performed RNA sequencing on laser-captured oculomotor neurons and spinal motor neurons from human post-mortem tissues and identified upregulated genes in oculomotor neurons relative to spinal motor neurons [30]. We took genes highly expressed in more than 5 resistant clusters identified earlier and converted them to human orthologs, we found there is a significant overlap between these genes and upregulated genes in oculomotor neurons (Hypergeometric test, P < 0.05, Figure 3G) while highly expressed genes in more than 5 vulnerable clusters do not show any significant overlap. This suggests that the transcriptional state of resistant neurons we identify is conserved.

If this conserved signature was indeed relevant to disease, we would assume that in end-stage disease there would be a mis-regulation of genes associated with this signature, as vulnerable neurons would be depleted whereas resistant neurons (where these genes would be upregulated) would be over-represented. We then turned to FTLD-TDP (frontotemporal lobar degeneration characterised by TDP-43 inclusions, also referred to as FTD) patient data, to check whether the conserved transcriptional signature identified above, composed of 97 genes, was indeed mis-regulated in end-stage disease. We looked to see if our genes of interest were altered in human cases, when using existing differential expression data [31]. When not adjusting for cellular composition, 20 fly resistant genes were transcriptome-wide differentially expressed comparing patients with a *C9orf72* repeat expansion to controls, and 42 comparing patients without a *C9orf72* repeat expansion to controls (Table S6A), the latter being significantly higher than expected based on the entire RNAseq dataset by Fisher exact test (p-value = 0.001). Notably, there was no significant overlap in any of the comparisons after accounting for differences in cellular composition (Tables S6B), suggesting that the differences seen are indeed related to differences in cellular composition. This suggests that the conserved signature modulating cell vulnerability is potentially relevant to disease. We next identified disease-associated modules of co-expressed genes that were enriched for biological processes (GO BP) or cellular components (GO CC) related to the endoplasmic reticulum and/or proteasome (Figure 3H), the pathways associated with resistant clusters in our fly model. In the analysis without adjustment for cellular composition, we found 4 disease-associated modules enriched for these pathways when comparing patients with a *C9orf72* repeat expansion to controls (Figure 3H, Table S7A), and 2 disease-associated modules when comparing patients without a *C9orf72* repeat expansion to controls (Table S7B). When adjusting for cell type proportions, we found 4 disease-associated modules enriched for these pathways when comparing patients with a *C9orf72* repeat expansion to controls (Table S7C), but no disease module was found when comparing patients without a *C9orf72* repeat expansion to control (Table S7D). This again suggests that pathways modulating vulnerability are mis-regulated in disease, although the signature itself does not appear to be specific to C9 and might be observed in patients without this expansion as well.

### *Xbp1* can modulate C9 toxicity

We next wanted to identify critical nodes in the gene or protein networks modulating vulnerability as these could be potential candidates for interventions to modulate disease.

To identify these, we placed all the proteostasis related genes in our starting point up in resistant and down in vulnerable DE analysis into a protein-protein interaction network and identified Regulatory particle triple-A ATPase 1 (*Rpt1*) as the central node with the highest connectivity (Figure 4A). *Rpt1* encodes an AAA-type ATPase which is part of the regulatory complex of the 26S proteasome [32, 33]. Remarkably, over-expression of *Rpt1* in all neurons increased the lifespan of C9 repeats expressing flies (Fig 4B), suggesting that increasing proteasomal activity across all neurons makes vulnerable neurons better able to deal with C9 toxicity.

**Figure 4.**
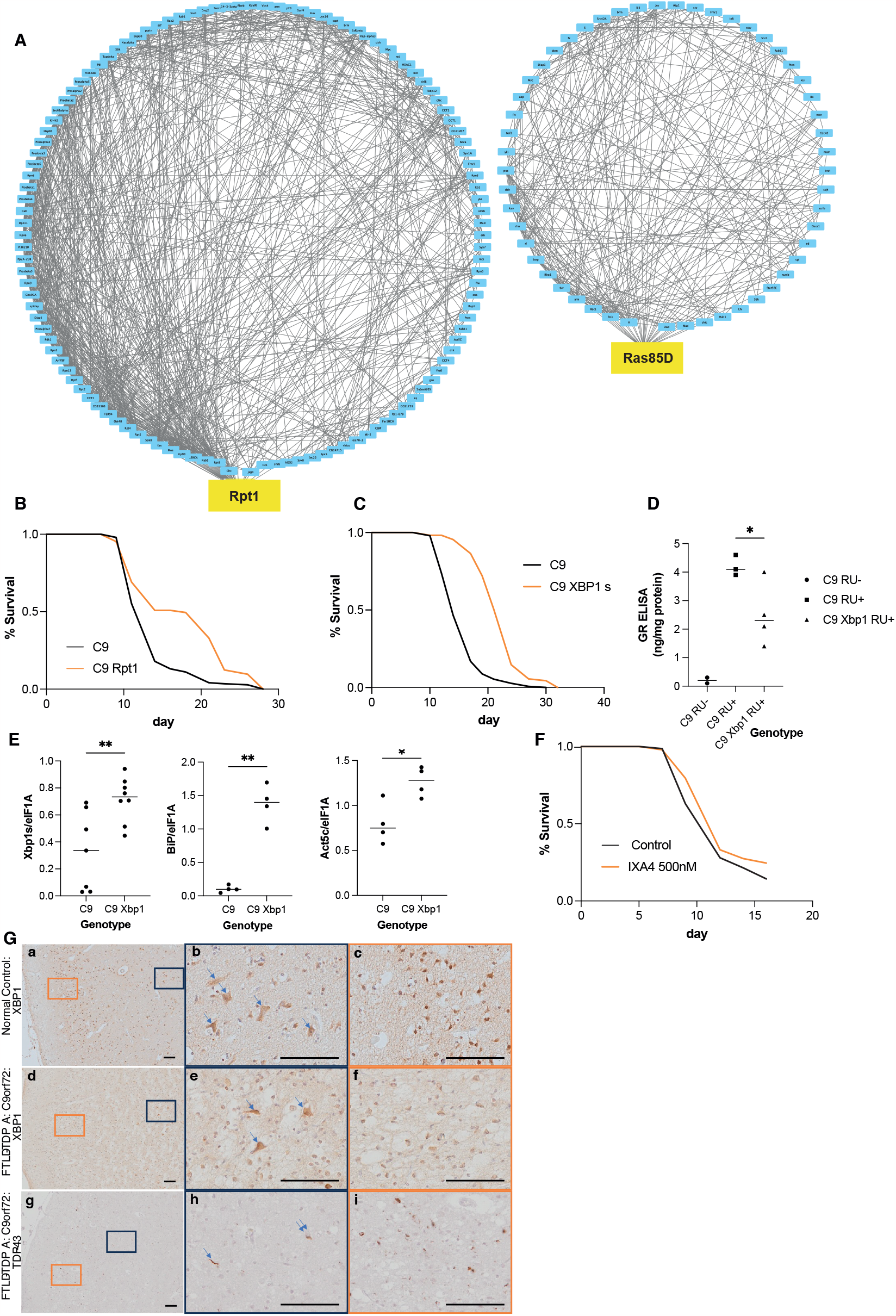
Validation of *Xbp1s* and *Rpt1* in C9 Fly Model. (A) Protein-protein interactions of genes that highly expressed in resistant clusters and lowly expressed in vulnerable clusters. (B) Lifespan of C9 fly model overexpressing *Rpt1* was significantly extended compared to C9 w1118 control (Log-rank test, p-value < 0.05). Genotype: w1118; UAS-36R; elavGS, w1118; UAS-36R; elavGS/UAS-Rpt1 (C) Lifespan of C9 repeats fly model overexpressing *Xbp1s* was significantly extended compared to C9 WDah control (Log-rank test, p-value < 0.05). Genotype: wDah; UAS-36R; elavGS, wDah; UAS-36R; elavGS/UAS-Xbp1s (D) GR ELISA of fly heads from the C9 repeats fly model uninduced (RU-) and induced (RU+) with and without *Xbp1s* co-expression (one-way ANOVA, post-hoc p-value < 0.05). (E) qPCR for Xbp1, BiP and Act5c on fly heads from the C9 fly model with and without *Xbp1s* over-expression. (F) Lifespan of the C9 repeats expressing fly model fed on control vehicle food and fed 500mM IXA4, an Xbp1s activator (Log-rank test, p-value < 0.05). (G) XBP1 (a-f) and TDP43 (g-i) immunohistochemistry on post-mortem human frontal cortex from a normal control and a FTLD-TDP type A case with a confirmed *C9orf72* expansion repeat. (a-c, arrows) mark XBP1 staining in the normal control. (e, arrows) mark strong XBP1 stain in the FTLD-TDP case. TDP43 neuronal cytoplasmic inclusions are marked in (h, double arrow) and those in neurites in (h, arrow). Bar = 100µm.

We also ran a transcription factor prediction algorithm iRegulon to predict the key TFs regulating the proteostasis gene network identified in our DE analysis, and the top TF identified was Xbp1 (Table S8), a well-known regulator of the Unfolded Protein Response, which has been implicated in Alzheimer’s disease [34-36].

To test whether Xbp1 could indeed modulate toxicity, we over-expressed an active form of *Xbp1* (spliced *Xbp1, Xbp1s*), together with C9 repeats and found that, indeed, Xbp1s over-expression led to a substantial lifespan increase of our C9 fly model (Figure 4C). We confirmed over-expression of *Xbp1s* by qPCR and showed it was leading to the increased transcription of known targets such as *BiP* (Fig 4E) and targets identified in our analysis, such as *Act5c*, which was the top DE Xbp1 regulated gene (Table S9), suggesting Xbp1*s* over-expression was indeed leading to the expected transcriptional response.

To further investigate how overexpressing of *Xbp1s* was leading to a rescue we performed GR ELISA on fly heads overexpressing C9 repeats with and without *Xbp1s*. The results showed that overexpressing *Xbp1s* significantly reduced the GR level in the fly head (Figure 4D), indicating that an increase in protein homeostasis across all neurons leads to a descrease of toxic peptides and an amelioration of disease phenotypes. To further validate the effect of Xbp1s activation on C9 repeats toxicity, we tested the lifespan of C9 repeats expressing flies fed a drug known to activate Xbp1s, IXA4. Feeding 500nM of IXA4 significantly extended the lifespan of C9 repeats expressing flies (Figure 4F) [37].

To check for disease relevance, we evaluated conserved *XBP1* targets in human cases’ brain transcriptomic analysis, we took genes over-expressed in resistant neurons and down-regulated in vulnerable neurons at baseline containing the *Xbp1* binding motifs based on iRegulon. Those genes were converted to human gene orthologues and in this way, we identified 40 human homologues to the *Xbp1* target genes, we compared these to DE genes from an FTLD-TDP human patient dataset [31] and identified 7 transcriptome-wide differentially expressed genes comparing patients with a *C9orf72* repeat expansion to controls and 13 comparing patients without a *C9orf72* repeat expansion to controls (Table S10A), the latter pairwise comparison is significantly higher than expected (p-value = 0.005). Notably, there was no significant overlap in any of the comparisons after accounting for differences in cellular composition (Tables S10B), again suggesting the difference is due to differential cellular composition in end-stage disease.

We next tested whether XBP1 is present in human neurons in disease relevant areas, such as the frontal cortex, by staining control and FTLD-TDP patient brains for XBP1 (Figure 4G). XBP1 shows strong neuronal staining in neurons throughout the cortical layers in control brains (Fig 4G a-c, arrows) whereas in the FTLD-TDP case there was a reduction in the number of neurons observed due to atrophy. However, the surviving neurons found in the deeper cortical layers in the FTLD-TDP case were positive with XBP1 (Fig 4G e, arrows), whereas the neurons in the upper cortical layers had a reduction in XBP1 staining. TDP-43 immunohistochemistry on sequential sections of the frontal cortex showed pathological inclusions in the form of neuronal cytoplasmic inclusions (Fig 4G h, double arrow) and neurites (Fig 4G h, arrow), confirming the presence of disease-associated pathology.

These experiments suggests that *Xbp1* drives a conserved transcriptional response, modulating neuronal vulnerability, and that *Xbp1* over-expression leads to an increase in neuronal resistance to C9 repeats toxicity.

## Discussion

In this study we look at the response to C9 repeat expression in a fly model of disease. We find that neuronal populations respond differentially to the expression of these repeats, with some degenerating quite rapidly and others displaying no apparent reduction in cell number, suggesting resistance to the toxicity associated with these repeats.

In particular we identify 17 neuronal clusters and 1 glial cluster as depleted upon C9 expression. The presence of a depleted glial cluster suggests that C9 repeats are able to induce cell death non autonomously in glial populations, given we only drove expression of the repeats in neurons, this suggests a potential role of glial degeneration in disease.

Our analysis concentrated on the 17 vulnerable neuronal populations. Interestingly, some of these populations (KC, Tm2, Tm4, Mi, T4/T5, for example) overlap to those found in a previous study using fly models of Tau and alpha-Synuclein toxicity [38], suggesting potentially common vulnerabilities across models of intracellular toxicity. This is also reflected in patients, where different pathologies often co-occur [39]. Other cell types appear uniquely vulnerable to C9 toxicity (T1, L2/L3), we do note however that the methodology for identifying vulnerable/resistant populations were identified in the two studies is different.

Some neurons (KC) degenerated very early in disease development, while others showed a later depletion (Tm3), this is similar to the patient setting, where cell death is initiated in one cell type and that this spreads, often along anatomical connections [6], in FTD/ALS this has been suggested to occur by disseminating TDP-43 pathology along cortical projections [40], it will be interesting to see if this is also occurring in our model.

Through differential expression analysis, we have identified pathways which are downregulated in vulnerable and resistant neurons. Vulnerable neurons uniquely downregulated the mitochondrial electron transport chain, mitochondrial related pathways were also implicated in toxicity in the single-cell sequencing analysis of Tau and alpha-synuclein fly models [38], suggesting common pathways across diseases.

Interestingly pathways related to ribosomes and translation were downregulated in both vulnerable and resistant neurons; unsurprisingly, this signature is often identified in bulk RNA sequencing [41] and the role of translation has been extensively described in C9 toxicity [25-27]. Our analysis reveals this as a common response across all neurons, not specific to the vulnerable ones, it is therefore possible that vulnerable neurons might be more susceptible to loss of translation.

Remarkably, our data suggests that the initial transcriptional state of neurons is a strong determinant of vulnerability. We find that protein homeostasis pathways are upregulated in resistant neurons and downregulated in vulnerable neurons, in control conditions, suggesting that resistant neurons are able to cope better with stress associated with toxic proteins. This is consistent with the fact that many FTD/ALS risk genes are involved in protein homeostasis pathways [42]. We also find that upregulation of key interactors or transcription factors, such as Rpt1 and Xbp1, can confer resistance to toxicity across all neuronal populations, indicating that promoting resistance pathways in vulnerable neurons can boost their resilience to toxicity.

Our study highlights, specifically, the role of the proteasome and the Unfolded Protein response in toxicity. Rpt1 is a key component of the 26S proteasome, which has been extensively implicated in disease development: pharmacological inhibition of the proteasome leads to the accumulation of TDP-43, SOD1 and poly-ubiquitinated proteins in cell culture models [43, 44], and mutations in proteins implicated in the ubiquitin-proteasome system, such as Ubiquilin-2 (*UBQLN2*) [45], *CCNF* [46] and *VCP* [47] can cause ALS/FTD. We show that upregulating a component of the proteasome is able to modulate toxicity in our fly model, suggesting it as a potential therapeutic target.

We also find that Xbp1 is a key transcription factor modulating the vulnerability of neurons to C9 toxicity. Xbp1 is predicted to control the expression of a large number of genes which are highly expressed in resistant and lowly expressed in vulnerable neurons and over-expression of the active form of Xbp1 is able to dramatically rescue C9 toxicity. *Xbp1* has been previously implicated in neurodegenerative disease, most notably AD, a polymorphism in *Xbp1* in humans has been potentially linked with Alzheimer’s disease development in a Chinese Han population [34] and over-expression of *Xbp1* an ameliorate fly and mice models of Alzheimer’s disease [35, 48].

The proteasome and Xbp1 potentially provide a link to the ageing process, proteasomal activity declines with age [49] and Xbp1 has been shown to ameliorate brain ageing in mice [50] and flies [51], suggesting that a decline in their function could be a contributing factor leading to disease development with advancing age.

In this work we concentrated on pathways modulating toxicity (up in resistant and down in vulnerable neurons), but there are pathways uniquely upregulated and uniquely downregulated in resistant neurons which warrant future analysis.

Overall, this study shows that neuronal vulnerability is strongly dependant on the transcriptional state of neurons at baseline, that resistant neurons have a higher resilience to proteostatic insults and that leveraging the properties of resistant neurons could be a way of boosting resistance in vulnerable neurons.

## Methods

### *Drosophila* models

For single-cell RNA sequencing, virgin female UAS-36R flies were crossed with male elavGS flies and the offspring females were used for single-cell library preparation. For neuronal vulnerability/resistance validation, UAS-mCD8::GFP was first combined to UAS-36R. Then the virgin females from the combined line were crossed with different split-Gal4 driver lines males and the offspring females were used for fluorescent imaging, while the offspring females from UAS-mCD8::GFP females crossing to different split-Gal4 driver lines males were used as controls. For lifespan experiments, UAS-Xbp1s and UAS-Rpt1 were backcrossed for six generations into WDah and w1118 separately. The virgin females of those were crossed to combined UAS-36R; elavGS males to generate offspring females for lifespan experiments, while WDah and w1118 virgin females were crossed to UAS36R; elavGS to generate females as controls. All flies were reared at 25°C on a 12:12 light: dark cycle at constant humidity and on standard sugar-yeast medium (15g/L agar, 50 g/L sugar, 100 g/L autolysed yeast, 100g/L nipagin and 2ml/L propionic acid). Adult-onset, neuron–specific expression of UAS constructs was achieved by feeding flies 24–48 hours after eclosion SYA medium supplemented with 200μM mifepristone (RU486) to induce transgene expression.

### Brain dissection and single-cell suspension preparation

25 females per group were dissected in Schneider’s *Drosophila* Medium on ice for less than 1.5 hr. The brains were collected in PBS. Then the brains were centrifuged at 2,000 rpm at 4°C for 5 minutes. The supernatant PBS was substituted with 50μL dispase (3mg/mL solution in PBS) and 75μL collagenase (100mg/mL solution in PBS). The brains were incubated with enzymes at 25°C in a 500 rpm shaker for 5 minutes. The enzyme solutions containing the brains were gently homogenised with flame-rounded tips for another 15 minutes. The dissociated cells were centrifuged at 2,000 rpm at 4°C for 5 minutes and washed with 1mL 0.04% BSA in PBS for 3 times and the cells finalised in 200μL 0.04% BSA in PBS. The cell suspensions were then passed through 10μm cell strainers and then checked for viability and counted for cell numbers by LUNA-FL Dual Fluorescence Cell Counter.

### 10X Genomics

Single-cell libraries were generated using Chromium Next GEM Single Cell 3’ GEM, Library & Gel Bead Kit v3.1 according to the manufacturer’s protocol. Briefly, about 13,000 cells were added to each channel with a targeted cell recovery estimate of 8,000 cells. After generation of nanoliter-scale Gel bead-in-EMulsions (GEMs), GEMs were reverse transcribed in a C1000 Touch Thermal Cycler (Bio Rad) programed at 53°C for 45 min, 85°C for 5 min, and hold at 4° C. After reverse transcription, single-cell droplets were broken and the single-strand cDNA was isolated and cleaned with Cleanup Mix containing DynaBeads (Thermo Fisher Scientific). cDNA was then amplified with a C1000 Touch Thermal Cycler programed at 98°C for 3 min, 12 cycles of (98°C for 15 s, 67°C for 20 s, 72°C for 1 min), 72°C for 1 min, and held at 4°C twice. Subsequently, the amplified cDNA was fragmented, end-repaired, A-tailed and index adaptor ligated, with SPRIselect Reagent Kit (Beckman Coulter) with cleanup in between steps. Post-ligation product was amplified with a C1000 Touch Thermal Cycler programed at 98°C for 45 s, 14 cycles of (98°C for 20 s, 54°C for 30 s, 72°C for 20 s), 72°C for 1 min, and hold at 4°C. The sequencing-ready library was cleaned up with SPRIselect beads.

### Fly brain imaging

Fly brains expressing GFP by split-Gal4 drivers were dissected in PBS on ice. The brains were then fixed in 4% paraformaldehyde in PBS for 15 minutes and washed 3 times with PBS for 10 minutes each time. The brains were mounted in Vectshield and visualised with a Zeiss Airyscan 880. Single confocal planes were taken every 5μm on z-stack and planes were stacked together for all images. All settings were kept the same within an experiment.

### Lifespan analysis

Male and female flies were allowed to mate and lay eggs for 24 hours on agar grape plates with yeast. The eggs were collected and seeded at standard density in 50ml bottles with SYA. After eclosion, flies were allowed to mate for 24–48 hours. At least 110–150 females of the appropriate genotype were split into groups of 15 and housed in vials containing SYA medium with or without drugs. Deaths were scored and flies tipped onto fresh food 3 times a week. Data are presented as cumulative survival curves, and survival rates were compared using log-rank tests. All lifespans were performed at 25°C.

### qPCR

Total RNA was extracted from 10-15 fly heads per sample using Trizol (Invitrogen) and subsequently treated with DNAse I (Ambion) for DNA digestion. The RNA was then reverse transcribed using Superscript II (Invitrogen) with oligo(dT) primers. Quantitative gene expression analysis was performed on a 7900HT real-time PCR system (Applied Biosystems) using SYBR-green technology (ABI). Relative quantities of transcripts were determined using the relative standard curve method normalized to eIF.

Primers used are:

Act5C_F: AAGTACCCCATTGAGCACGG

Act5C_R: ACATACATGGCGGGTGTGTT

BiP_F: TCTTGTACACACCAACGCAGG

Bip_R: CAAGGAGCTGGGCACAGTGA

eIF-1A_F: ATCAGCTCCGAGGATGACGC

eIF-1A_R: GCCGAGACAGACGTTCCAGA

Rpt1_F: CTTATGGGCAGAGCCAGTATCA

Rpt1_R: GATTCCAGTCAGCTCGTTCAC

Xbp1s_F: CCGAACTGAAGCAGCAACAGC

Xbp1s_R: GTATACCCTGCGGCAGATCC

### GR ELISA

10 female frozen heads per tube were homogenised in 2% SDS buffer containing 1X RIPA buffer and complete mini EDTA-free protease inhibitor cocktail at Room Temperature for about 30 sec or until the heads were no longer intact. Samples were then heated at 95C for 10 min. After centrifuged at 14000 rpm for 20 min at RT, the supernatants were collected in the new tubes and stored at −80°C. The protein concentration was determined using Pierce BCA Protein Assay Kit according to the manufacturer’s manual. Samples were diluted to the same concentration with homogenisation buffer and 25μL were loaded in duplicate in the 96-well Meso Scale Discovery (MSD) immunoassay plate. The singleplex MSD immunoassay was performed to measured poly-GR expression levels as described before [26].

### Human Briain Staining

Brain sections were first prepared as follows: Initially, sections were selected at the desired level and baked at 60°C. Subsequently, deparaffinization was carried out with two rounds of 100% xylene for 5 minutes each, followed by four immersions in 100% ethanol for 2 minutes each. Endogenous peroxidase activity was inhibited by incubating the sections for 10 minutes in a 3% H2O2 solution in dH2O. Antigen retrieval involved preparing a buffer, boiling it in a pressure cooker for 14 minutes with the lid off, and then placing the slides in the buffer with the lid on, cooking for an additional 5 minutes at high heat. After antigen retrieval, slides were washed with water. To prevent non-specific antibody binding, PAP pen rings were applied around the tissue, and a 1% BSA blocking solution was added to the sections for 10 minutes. Primary antibodies (ab37152 for Xbp1) were applied and incubated for 2 hours to overnight at room temperature. Following primary antibody incubation, the sections were washed in 1xTBS for 10 minutes, twice. Subsequently, biotinylated secondary antibodies were applied for 1 hour. Avidin-biotin complex (ABC) was prepared and applied for 30 minutes. After each incubation step, the sections were washed in 1xTBS for 10 minutes. DAB (3,3’-Diaminobenzidine) solution was used to develop the signal, followed by thorough rinsing in water. Sections were then dried, cleared in 100% Histoclear (or xylene), coverslipped with DPX or DePeX mounting medium, and left to dry overnight. To remove excess DPX, ethanol was applied and gently scraped off with a razor blade. Finally, the prepared slides were observed under a light microscope.

## Quantification and statistical analysis

### 10X Data Preprocessing

The raw sequencing data was aligned to the FlyBase Drosophila melanogaster (r6.43) by CellRanger (6.1.2) with a customization of adding the sequence of GeneSwitch to the genome. The generated matrices were first filtered out cells with less than 200 genes and genes detected less than 3 cells. We also filtered out cells with more than 3000 genes and cells with mitochondria DNA higher than 5%. The data was then run through MNN batch correction among 3 batches, and after visualising by UMAP, we decided to remove one batch in day8 group as it looks off on UMAP. The rest 101,806 high-quality cells were taken to the further analysis.

### Clustering and visualisation

Following the standard scanpy workflow, all gene counts from each cell were normalised by the total counts of the cell and log1p-transformed. 902 highly variable genes were determined by scanpy and used for PCA analysis. We then processed with 40 PCs and k value of 20 for visualisation with UMAP. Clustering was using leiden with resolution at 10.0, which assigned all the cells into 214 clusters.

### Cell type annotation

Broad cell-type annotations of each cluster were inferred from classifier derived cell-type labels as well as neuronal (*elav, nSyb*), glia (*repo*), photoreceptor (*chp*) and hemocyte (*Hml*) marker genes. Neurotransmitter identities were assigned based on previously established neuron-neurotransmitter connections as well as Acetylcholine (*VAChT, ChAT*), Glutamate (*VGlut*), GABA (*Gad1*), Dopamine (*ple*), serotoninergic (*SerT*), Tyramine/Octopamine (*Tdc2*) and Histamine (*Hdc*) metabolism marker genes.

More specific cell type annotation was performed with PopularVote by transferring the labels from reference fly brain atlas to our dataset. The leiden clusters which have agreement over 70% with PopularVote were assigned with cell type labels from fly brain atlas.

### Differential abundance analysis and determination of vulnerable/resistant clusters

Cell number change across time points was calculated with milopy. To achieve a roughly evenly distributed cell numbers in each neighbourhood, parameter of proportion was set to 0.1. The FDR for significantly changed neighbourhood was set to 0.1. The cell number change in each cluster was calculated in the following formula.

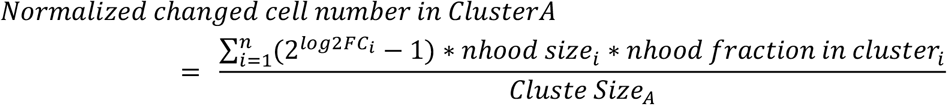

Depending on the cell number changes, all the clusters were divided into four categories – clusters with normalised changing cell number below −0.2 as depleted clusters, clusters with normalised changing cell number over 0.2 as increasing clusters, the rest of clusters as resistant clusters. In the resistant clusters, we assigned some clusters as moving clusters because they have over 20% of cells being increased and decreased at same time and we think the transcriptome profiles in the cluster were also changed due to the C9 toxicity but not enough to make the cluster being depleted.

### Differential expression analysis and Gene Ontology pathway analysis

Differential expression analysis was performed with scanpy’s built-in “find_marker_genes” with Wilcoxon rank test and the results were filtered with adjusted p values of 0.05. The differentially expressed genes were run through GOATOOLS for pathway analysis using Fisher’s exact test as enrichment analysis and the p values were corrected by Benjamini-Hochberg procedure, and 0.05 was set to the threshold of significance.

### Human RNAseq Data

We leveraged published human frontal cortex RNA sequencing (RNAseq) data that included frontotemporal lobar degeneration (FTLD) patients with or without motor neuron disease, who either have (n=34; C9Plus) or do not have a *C9orf72* repeat expansion (n=44; C9Minus), as well as control subjects without any neurological disease (n=24; Control) (Dickson DW, et al: Extensive transcriptomic study emphasizes importance of vesicular transport in C9orf72 expansion carriers. Acta Neuropathol Commun 2019, 7(1):150.). Pairwise comparisons were previously completed for differential expression and weighted gene co-expression network analyses (WGCNA), both with and without adjustment for cellular composition. To determine whether differential expression results were enriched for genes of interest, we utilized a Fisher’s exact test.

## Data availability

The scRNA-seq raw data has been deposited here: EBI: E-MTAB-12440

## Acknowledgements

We thank Dr Dune and Dr Avsec for their advice regarding the single-cell sequencing analysis. We thank Nazif lab from IHA UCL for providing the Xbp1 fly line. We also thank Sanbomics from Youtube for posting informative videos of single-cell RNA sequencing analysis tutorials. This work was supported by the ARUK Senior Fellowship ARUK-SRF2018A-003 (TN), an MRC grant MR/V003585/1 (TN), a Dunhill Studentship (TN) and a Malaysian Government Studentship (SA).

## Notes

### Competing Interest Statement

The authors have declared no competing interest.

### Summary of Updates

Revised to a higher quality pdf with the figure being more clear

